# Light-entrained chromatin priming poises rapid metamorphosis in a marine sponge

**DOI:** 10.64898/2026.01.28.702437

**Authors:** Huifang Yuan, Oceane Blard, Zac Pujic, Bernard M. Degnan, Sandie M. Degnan

## Abstract

The most widespread animal life cycle includes a planktonic larval stage that is environmentally induced to settle and rapidly metamorphose into a benthic juvenile. Although this transition is critical for survival, how planetary cues such as light interact with local ecological signals to prepare the genome for such rapid reprogramming remains poorly understood. Here, using the sponge *Amphimedon queenslandica*, we integrate time-resolved transcriptomic and chromatin accessibility profiling across larval competence, settlement, and the first hours of metamorphosis. We show that diminishing light at sunset is associated with extensive chromatin remodelling in swimming larvae prior to settlement, despite relatively modest changes in gene expression. Upon environmental induction by a coralline alga, metamorphosis is accompanied by the rapid and transient activation of a large suite of deeply conserved transcription factors, including AP-1/bZIP family members, whose binding motifs are enriched in newly accessible regulatory regions near differentially expressed genes. Larvae prevented from experiencing sunset by constant light fail to settle and instead adopt an alternative transcriptional and chromatin state characterised by widespread loss of accessibility and repression of competence-associated transcription factors, including the circadian regulator CLOCK and other bHLH-PAS factors. These observations support a model in which light-entrained transcriptional activity establishes an anticipatory, permissive chromatin landscape that enables immediate and coordinated transcriptional reprogramming upon environmental induction. We propose that such chromatin priming may represent a broadly deployed regulatory strategy underlying the speed and robustness of metamorphosis in biphasic animal life cycles.

## Introduction

The biphasic pelagobenthic life cycle is widespread in early-diverging animal lineages (sponges and cnidarians) and in molluscs, echinoderms, annelids and several other bilaterian phyla, suggesting it evolved before metazoan cladogenesis (Degnan and Degnan, 2010, 2006; Hadfield, 2000; Hedgpeth and Jagersten, 1974; Martín-Zamora et al., 2023; Nielsen, 1998; Rieger, 1994). This ancient aquatic life cycle begins with endogenously-regulated embryogenesis that produces a ciliated planktonic larva, and culminates with settlement and metamorphosis into a benthic juvenile. In contrast to embryogenesis, metamorphosis is characterised by three defining features: environmental induction, coordinated transformation of one complex body plan into another, and a dramatic ecological transition (Hadfield, 2000; Hedgpeth and Jagersten, 1974; Nielsen, 1998). Despite the scale of morphological and physiological change required, in diverse marine invertebrates this transition occurs strikingly quickly as a means to enhance post-settlement survival (Conaco et al., 2012; Hadfield, 2000; Heyland and Moroz, 2006; Wray, 1999).

Marine larvae are strongly influenced by planetary constants, including daily, tidal, lunar and seasonal rhythms, with gametes or larvae being released at a species-specific time. Most larvae also have developmentally-regulated behaviours that enable settlement to be induced by species-specific biochemical cues that are associated with a favourable settlement site for adult survival and reproduction (Bishop et al., 2006a; Ettinger-Epstein et al., 2008; Hadfield et al., 2001; Hodin et al., 2015; Randall et al., 2024). The capacity to respond to these cues – widely called “larval competence” – develops while larvae swim in the plankton over timescales ranging from minutes to weeks, depending on the species (Bishop et al., 2006b; Hadfield, 2000; Hodin et al., 2015; Jackson et al., 2002; Randall et al., 2024; Say and Degnan, 2020). Together, these traits indicate that larvae across disparate animal phyla have evolved to integrate constant planetary signals with local ecological cues to optimise dispersal, settlement success and subsequent survival.

Despite the prevalence and ecological importance of this life cycle, it remains unclear how the metazoan genome integrates planetary and ecological signals to regulate the acquisition of larval competence, the decision to settle, and the rapid remodelling of the body plan at the onset of metamorphosis. Here, we address this problem by exploiting the experimental tractability of the sponge *Amphimedon queenslandica*, in which larval release, competence, settlement and metamorphosis are well characterised and readily manipulated (Blard et al., 2026; Conaco et al., 2012; Degnan et al., 2015; Degnan and Degnan, 2010; Jindrich et al., 2017; Nakanishi et al., 2015; Say and Degnan, 2020; Sogabe et al., 2016; Ueda et al., 2016; Wong et al., 2022). We experimentally induce competent larvae to settle on a naturally co-occurring coralline alga under natural and perturbed light conditions and profile chromatin accessibility and gene expression in the same individuals during the first hours of metamorphosis. This approach allows us to reveal that larvae anticipate impending settlement through diurnal light cues and the deployment of pioneer transcription factors that establish a permissive chromatin landscape prior to environmental induction. Given that sponges diverged from other animal lineages more than 700 million years ago, elucidating these mechanisms provides insight into how deeply conserved gene regulatory programs enabled the evolution of rapid, environmentally induced metamorphosis in the ancestral biphasic animal life cycle.

## Results

### Dramatic changes in gene expression in the first hour of metamorphosis

*Amphimedon queenslandica* larvae emerge from adult sponges in the early afternoon and become competent to settle on the coralline alga *Amphiroa fragilissima* 5-6 hours later, shortly after sunset (Degnan and Degnan, 2010; Say and Degnan, 2020). We have shown previously that diminishing light at sunset is necessary for larvae to respond to the algal cue under normal conditions and to initiate settlement and metamorphosis (Say and Degnan, 2020). The first hour following settlement is characterised by extensive cellular reprogramming and the rapid dissolution of the larval swimming anteroposterior axis (Figure 1A-C; Figure 1 – video 1) (Blard et al., 2026; Degnan et al., 2015; Leys and Degnan, 2002; Nakanishi et al., 2015, 2014; Sogabe et al., 2016).

**Figure 1.**
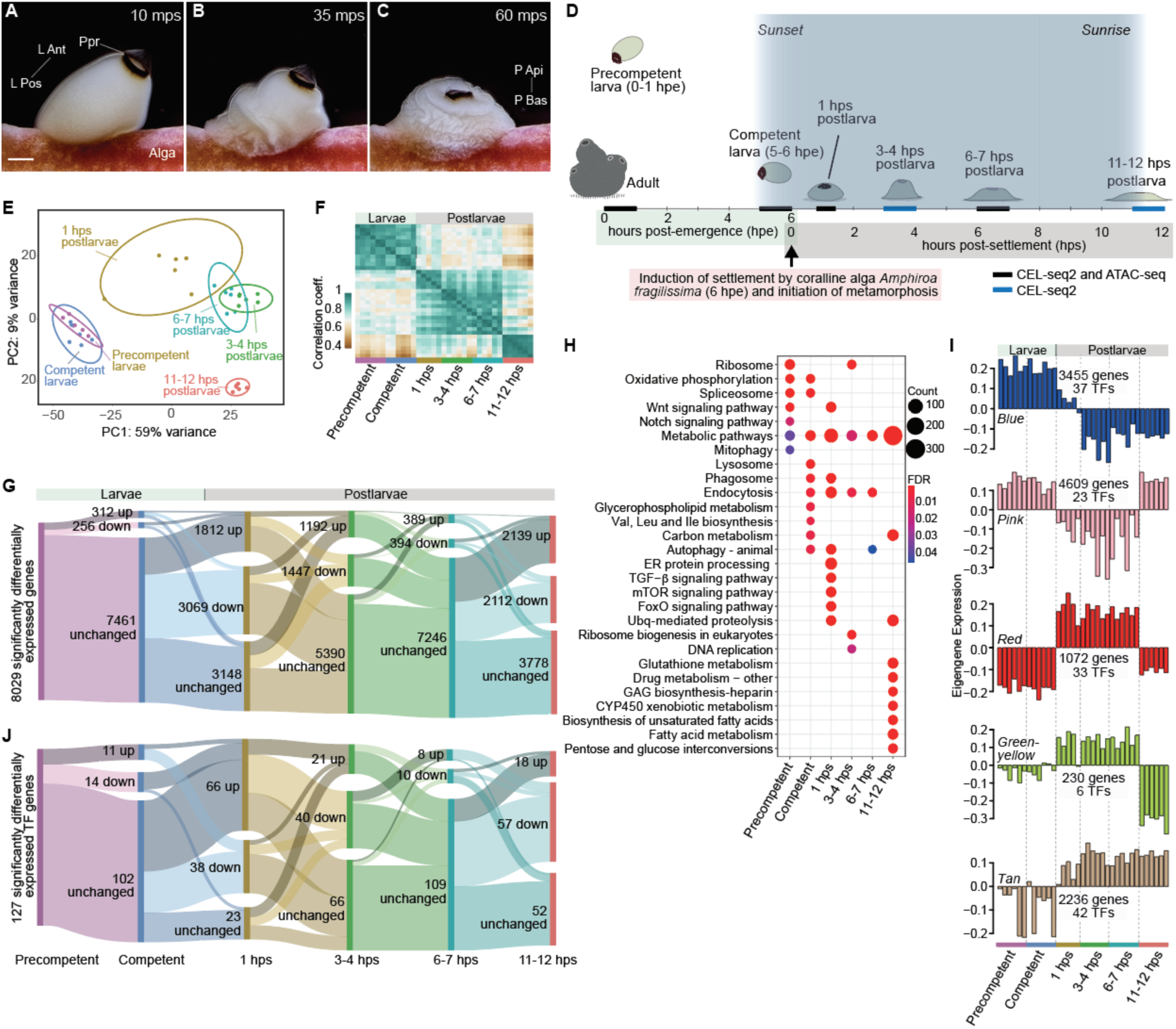
Gene expression during larval development and the initiation of sponge metamorphosis. A-C, Photomicrographs of the first hour of metamorphosis after competent *A. queenslandica* larvae settle on the coralline alga *A. fragilissima* (Alga). L Ant and L Pos, larval anterior-posterior axis; P Api and P Bas, postlarval apical-basal axis; Ppr, posterior pigment ring; mps, minutes post-settlement; scale bar, 100 µm (see Figure 1 - video 1) (Blard et al., 2026). **D,** Timeline of larval and early postlarval developmental stages analysed using CEL-seq2 and ATAC- seq (see Methods). Larvae become competent to respond to an inductive cue associated with *A. fragilissima* just after sunset, 4-6 h after emerging from the adult sponge (Degnan and Degnan, 2010; Say and Degnan, 2020). **E,** Principal component analysis (PCA) of CEL-seq2 transcriptomes with 95% confidence level ellipses shown; n = 6 for each stage. **F,** Hierarchical clustered heatmap of Pearson correlation coefficients of replicated larval and postlarval transcriptomes based on DESeq2-normalised counts of the 8,029 significantly differentially expressed genes (DESeq2; *p*-adj < 0.1). **G,** Alluvial plot showing dynamics of differentially expressed genes through larval development and early metamorphosis. **H,** Top 10 significantly enriched KEGG pathways (FDR < 0.05) based on significantly upregulated genes at each developmental stage; GO-MWU analysis of upregulated genes reveal stage-specific enrichments largely consistent with the KEGG analysis (Figure 1 – figure supplement 1; Supplementary file 4). FDR, false discovery rate. **I,** WGCNA co- expression modules that comprise genes that are down- (blue, pink) and up- (red, yellow-green, tan) regulated at the start of metamorphosis. Number of coding genes and TFs are shown. **J,** Alluvial plot showing dynamics of 127 significantly differentially expressed TF genes as per Figure 1G.

To investigate transcriptional and chromatin changes associated with the acquisition of competence and the onset of metamorphosis at sunset, we analysed biologically replicated transcriptomes from precompetent larvae, competent larvae, and postlarvae at 1, 3-4, 6-7 and 11- 12 hours post-settlement (hps) (see Methods; Figure 1D; Supplementary file 1). Across this time course, 8,029 genes were differentially expressed (DESeq2; *p*-adj < 0.1). Swimming larvae, early postlarvae (1, 3-4 and 6-7 hps) and later postlarvae (11-12 hps) showed transcriptionally distinct states (Figure 1E-G; Supplementary files 2 and 3).

The most pronounced transcriptional shift occurred within the first hour of metamorphosis, with 4,881 genes differentially expressed from competent larval to 1 hps postlarval stages; of these, 63% were downregulated (Figure 1G; Supplementary file 3). These repressed genes were enriched for diverse metabolic processes (Figure 1H; Supplementary file 4). In contrast, genes transiently upregulated during this interval were involved in Wnt, TGF-β, mTOR and FoxO signalling, and endocytosis (Figure 1H; Supplementary file 4). Together, these results indicate a rapid shift from a larval physiological state to a metamorphic developmental program, coincident with the onset of widespread reprogramming of larval cell types and reorganisation of the body plan (Blard et al., 2026). By 11-12 hps, gene expression shifted back towards metabolic processes, reflecting the establishment of a new postlarval physiological state (Figure 1H; Supplementary file 4).

Weighted gene co-expression network analysis (WGCNA) assigned 69% of expressed genes to five co-expression modules, typified by either up- or downregulation between competent larval and 1 hps postlarval stages (Figure 1I; Figure 1 – figure supplement 1; Supplementary file 5). These modules reinforce the conclusion drawn from the DE-Seq2 analysis that the first hour after settlement represents the most transcriptionally dynamic phase, marked by coordinated repression of larval metabolic programs and activation of developmental signalling pathways (Figure 1 – figure supplement 1).

### Conserved transcription factors are expressed at exceptionally high levels during metamorphosis

The sharp transcriptional reorganisation that characterises the first hour of metamorphosis is accompanied by a striking and selective induction of transcription factors (TFs). In contrast to the overall downregulation of gene expression at this stage, 63% of the TFs are significantly upregulated, suggesting roles in both gene activation and repression. Surprisingly, 52 of the 127 differentially expressed TFs are at levels higher than at least 95% of all expressed genes (Figure 2A, B; Supplementary files 3 and 6). These highly expressed TFs span multiple conserved families, including bZIP, bHLH, Tbox, Fox, Sp/KLF, homeobox and Sox.

**Figure 2.**
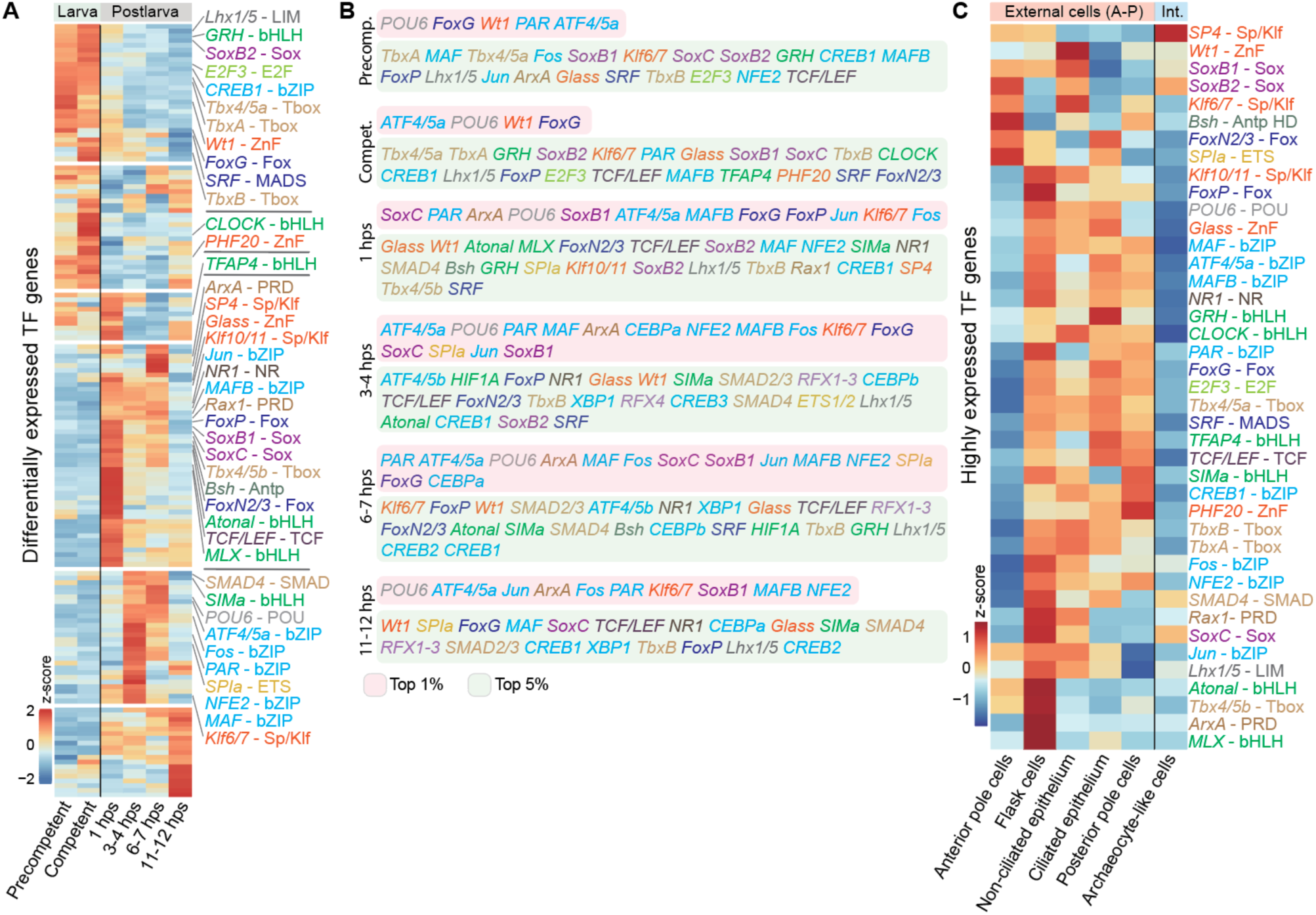
Dynamic and high-level expression of transcription factor genes during larval development and early metamorphosis. A, Scaled heatmap of 159 TF genes expressed during larval and postlarval development, of which 127 are significantly differentially expressed between at least two successive stages. The 40 most highly expressed TFs, all of which are among the top 5% of the most-highly expressed genes, are annotated to the right of the heatmap. TF genes names are followed by their gene family name, which is colour-coded into families; this colour- coding is used in all figures. **B,** Lists of TF genes in the order of their level of expression for all larval and postlarval stages. TFs in the pink and green boxes are amongst the top 1 and 5% of all expressed genes, respectively (Supplementary file 6). **C,** Scaled heatmap of the 40 most highly expressed TFs in larval cell types (Sebé-Pedrós et al., 2018). External epithelial cells are to the left - cell types ordered left-to-right based on their enrichment level along the anterior-posterior (A-P) axis; internal archaeocyte-like cells are to the right (Supplementary file 7).

All but one of these highly expressed TFs (the exception being the constitutively expressed SRF) share three features: (i) they are differentially expressed either during the acquisition of larval competency or within the first hour of metamorphosis; (ii) they belong to WGCNA modules that change sharply at the onset of metamorphosis; and (iii) they are significantly downregulated by 11-12 hps (Supplementary file 5). This coordinated temporal pattern is consistent with a transient but potent regulatory phase commencing at settlement and associated with large-scale reprogramming of larval cell states.

As larvae acquire competence to settle at sunset, 11 TFs are upregulated and 14 are downregulated (Figures 1J and 2A, B; Supplementary file 6). Among the upregulated and highly expressed TFs are two bHLH factors (*CLOCK* and *HIF-1*), the bZIP *ATF4/5*, two Fox TFs (*FoxN2/3* and sponge-specific *Fox 2*), *TboxB* and the zinc-finger (ZnF) TF *Glass*. *CLOCK* exhibits the most pronounced transient expression at competence, increasing from pre-competence expression (p- adj = 5.2e-6) and decreasing at 1 hps (p-adj = 1.5e-15) (Figure 2A; Supplementary file 6). In parallel, a sponge nuclear receptor (*NR1*) and multiple bZIP factors are significantly downregulated, including the highly-expressed AP-1 components *Fos* and *Jun*, and several of their potential partners (*CEBP*, *CREB*, *PAR*, *MAF* and *NFE2*) (Jindrich et al., 2017; Reinke et al., 2013;

Rodríguez-Martínez et al., 2017). These TFs, together with most other highly-expressed TFs, are enriched in externally facing larval cell types – ciliated and non-ciliated epithelia, and flask cells – that are implicated in detecting the algal inductive cue and that undergo rapid morphological and functional transformations during the first hour of metamorphosis (Figure 2C; Figure 1 – figure supplement 1; and Supplementary file 7) (Blard et al., 2026; Leys and Degnan, 2002; Nakanishi et al., 2014, 2015; Sebé-Pedrós et al., 2018a; Sogabe et al., 2016; Ueda et al., 2016).

In contrast to the widespread repression of gene expression that dominates the first hour of metamorphosis, the majority (63%) of differentially expressed TFs are upregulated during this period. Of these, 48 rank within the top 5% of all expressed genes (Figure 2A, B; Supplementary file 6). Notably, AP-1 and the bZIP factors that are downregulated at competence are now rapidly and strongly induced (Figure 2A; Supplementary file 6). In addition, TFs with deeply conserved roles in animal development (multiple Sox, bHLH, Fox, SMAD, Tbox and homeobox members, and NR1) and innate immunity (NF-κB, STAT and IRF) are transiently upregulated at the onset of metamorphosis (Figure 2A, B; Supplementary file 6). The rapid induction and exceptionally high expression of these TFs are consistent with a central role in coordinating environmentally-induced cell state transitions and establishing a robust regulatory environment capable of driving the rapid transformation of the larval body plan into that of the filter-feeding juvenile.

Similar to the dynamics observed for developmental signalling pathways, nearly all of these TFs are transiently upregulated during the first hour of metamorphosis and subsequently downregulated between 6–7 and 11–12 hps (Figure 2A, B; Supplementary file 6). Together, the scale, synchrony and tempo of TF activation and repression are consistent with the genome of competent larvae being poised to respond immediately to the algal inductive cue, enabling a rapid and coordinated transcriptional response that underpins the extensive cell reprogramming that transforms the larval body plan.

### Open chromatin regions are enriched in binding motifs of highly expressed transcription factors

To determine how the *A. queenslandica* genome responds to the inductive algal cue, we profiled chromatin accessibility in larvae and postlarvae using an assay for transposase-accessible chromatin with high-throughput sequencing (ATAC-seq) (see Methods; Figure 1D) (Buenrostro et al., 2015). We identified 60,716 non-overlapping open chromatin regions (OCRs; ATAC-seq peaks) that were highly reproducible across biological replicates and that overlapped with previously defined larval promoters and enhancers identified by ChIP-seq (Gaiti et al., 2017) (Figure 2 – figure supplement 1; Supplementary files 8 and 9). Consistent with patterns observed in *A. queenslandica* embryos and adults, as well as other animals (Cornejo-Páramo et al., 2022; Wong et al., 2020), approximately 80% of OCRs are located within protein-coding gene bodies or within 1 kb of transcription start or end sites.

Between 67% and 86% of differentially expressed protein-coding genes and genes belonging to the five WGCNA co-expression modules have proximal OCRs, depending on the stage or module. (Figure 3A, B; Figure 3 – figure supplement 1; Supplementary files 10 and 11). There are approximate three times as many expressed genes that have OCRs compared to genes that are not expressed (Figure 3A). Expressed genes without nearby OCRs may be regulated by more distal cis-regulatory elements or, given the high gene density of the *A. queenslandica* genome (Fernandez-Valverde and Degnan, 2016), by OCRs associated with neighbouring genes.

**Figure 3.**
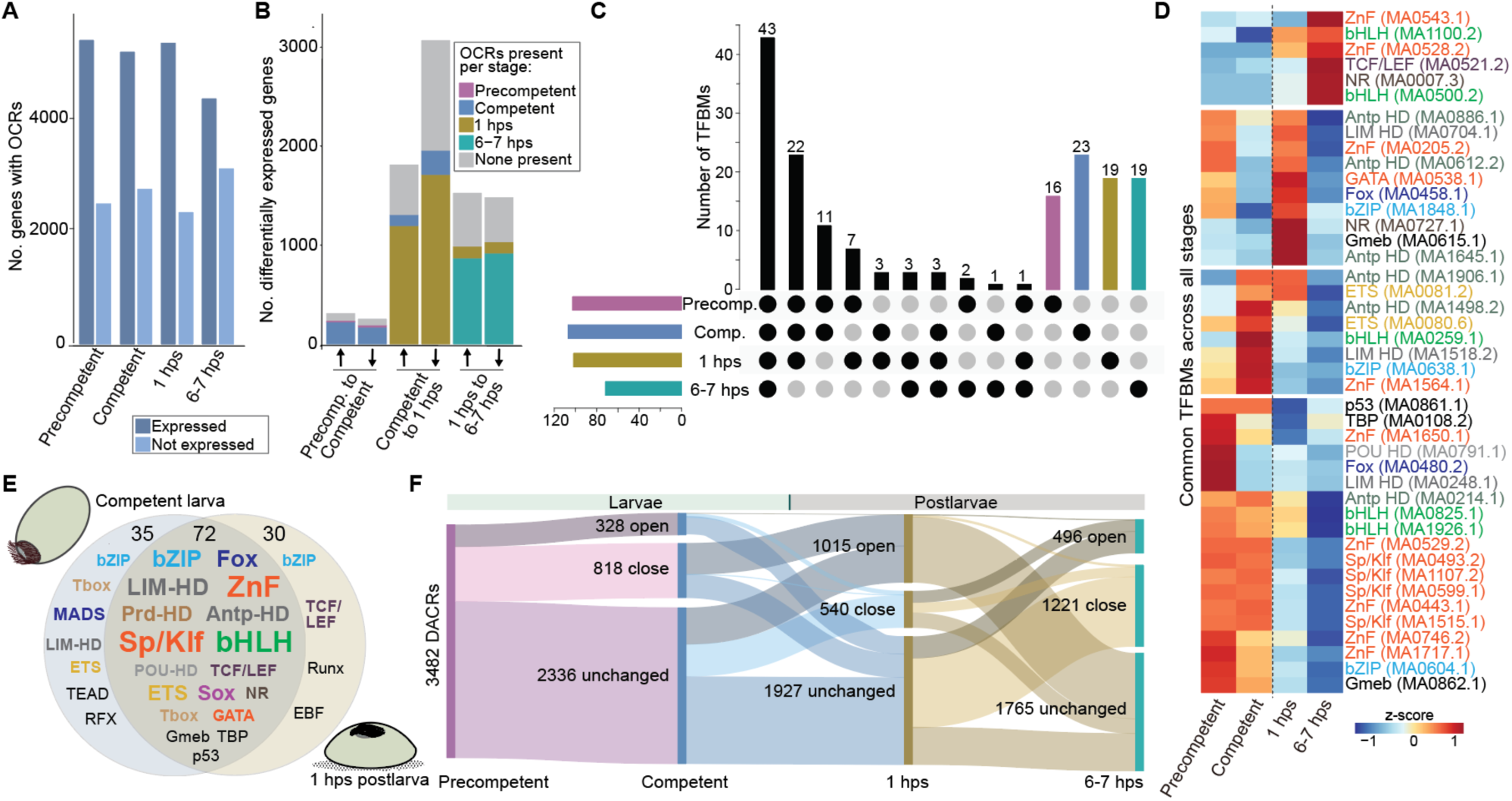
Open chromatin regions associated with expressed genes are enriched for binding sites of highly-expressed TFs. A, The number of OCRs associated with protein-coding genes expressed in larval and postlarval stages compared to genes not expressed at these stages. On average, there are 16,469 and 26,177 genes expressed and not expressed at these four stages, respectively (Supplementary file 10). **B,** The number of significantly up (↑) and down (↓) regulated genes that have nearby OCRs. Colour coding within the bars show at which stage the OCRs first appear. **C,** Upset plot showing the distribution of the 173 TFBMs shared and unique to each larval and postlarval stage. **D,** Heatmap of the 43 TFBMs that are significantly enriched in all larval and postlarval stages. TFBMs that potentially interact with highly expressed TFs are colour-coded by TF class as per Figure 2A; black, TFBMs of other expressed TFs. Jasper code follows TF class/family name to the right. **E,** Venn diagram of the differential enhancement of TFBMs in OCRs associated with genes that are differentially expressed between competent larvae and 1 hps postlarvae, with the TFBM class and family name size scaled to prevalence (Supplementary file 12). **F,** Alluvial plot showing DACRs across larval development and early metamorphosis. Open and close, chromatin accessibility significantly increases and decreases, respectively.

Previous studies have shown that sponge TFs can interact with bilaterian *cis*-regulatory elements and that sponge enhancers can drive developmental gene expression in vertebrates (Elliott et al., 2025; Richards et al., 2008; Wong et al., 2020). However, to avoid bilaterian-specific annotation biases – given that sponges and bilaterians diverged 700 million years ago – we restricted motif annotation to TF classes and families rather than individual TFs (see Methods). This analysis identified 173 TF binding motifs (TFBMs) corresponding to TF classes and families that are significantly enriched in OCRs associated with genes expressed during larval development and early metamorphosis (Homer p-value < 1e^-10^; Figure 3C; Supplementary file 10). Of these motifs, 78% correspond to predicted targets of the highly expressed TFs identified above, with an additional 15.6% attributable to highly expressed non-Sp/Klf ZnF TFs (93.6% total) (Figures 2A, B and 3D, E). These include 35 variants of the bHLH E-box motif (CACGTG), 40 homeodomain motifs (including POU, LIM and Antp), 17 Fox, 14 Sp/Klf and 9 bZIP motifs. The enrichment of these 173 TFBMs varies dynamically across developmental stages and co-expression modules, consistent with stage-specific changes in chromatin accessibility during larval development and metamorphosis (Figure 3D-F; Figure 3 – figure supplement 1; Supplementary files 10 and 11).

Across larval development and early metamorphosis, 3,482 chromatin regions exhibit significant changes in accessibility (differentially accessible chromatin regions; DACRs). More than 56% of differentially expressed genes have OCRs at the same developmental stage in which they significantly change expression, while 4-8% have accessibility one stage prior to the corresponding change in gene expression (FDR < 0.05; Figure 3B, F; Figure 3 – figure supplement 1; Supplementary files 3, 9 and 12). Notably, nearly one-third of DACRs (1,146) change accessibility between pre-competent and competent larvae, with 71.4% becoming less accessible at competence. This extensive chromatin remodelling contrasts sharply with the relatively modest number of transcriptional changes between these stages (Figure 1G), and suggests that many chromatin changes precede settlement and may contribute to establishing a permissive genomic state for the rapid transcriptional activation required at the onset of metamorphosis.

Consistent with this interpretation, 62% of all genes that are differentially expressed in 1 hps postlarvae (3032) have proximal chromatin regions already accessible in competent larvae (Supplementary files 3 and 9), although statistically we find no significant difference in the rate of differential expression between genes associated with newly opened OCRs and those associated with other OCRs (9.84% vs 11.47%; χ²(1) = 1.76, p = 0.184). Nonethless, during the first hour of metamorphosis, 65% of the 1,555 DACRs become more accessible, a proportion closely matching the fraction of genes that are transcriptionally repressed at this stage (63%; cf. Figures 1G and 2F). This correspondence is consistent with the involvement of rapidly induced, highly expressed TFs in transcriptional repression. It supports a model in which pioneer-like and other highly expressed TFs regulate both the acquisition of competence and the initiation of metamorphosis by interacting with their cognate binding motifs in OCRs proximal to differentially expressed genes.

### *CLOCK*, *Jun* and *Fos* OCRs are enriched in binding motifs of highly expressed TFs

Given a multitude of highly expressed TFs are also differentially expressed either at competence or at 1 hps and that more than 90% of TFBMs enriched in larval and postlarval OCRs are potential binding sites of these highly expressed TFs (Figures 2A, B and 3D, E; Supplementary files 6 and 10), we chose to analyse the OCRs associated with three TF genes that are differentially expressed and appear central to the acquisition of larval competence or the initiation of metamorphosis: *CLOCK*, *Jun* and *Fos*. bHLH and bZIP TFBMs are abundant in OCRs associated with genes that remain constantly expressed across these stages (Supplementary file 10), reinforcing the view that CLOCK and AP-1 occupy central positions in the regulatory networks governing both competence acquisition and environmentally induced cellular reprogramming at the onset of metamorphosis.

We analysed the *CLOCK* gene because it (i) is the most significantly and transiently upregulated TF gene in competent larvae at sunset, consistent with a light-dependent role in establishing competence, and (ii) is a pioneer TF that can be affected by environmental light in other animals (Figure 2A; Supplementary file 6) (Doi et al., 2006; Iwafuchi-Doi and Zaret, 2016; Jindrich et al., 2017; Menet et al., 2014). bHLH-PAS and bHLH (E-box) family motifs consistent with CLOCK:BMAL/CLK-like activity are present in OCRs associated with 165 differentially expressed genes at competence and with 1,121 at 1 hps (Supplementary file 13). This widespread distribution of bHLH-PAS/E-box family motifs is consistent with pioneer-like regulatory activity associated with CLOCK-related programs that may contribute to establishing a permissive genomic state for rapid transcriptional responses to the algal inductive cue.

In a genomic context, the *CLOCK* gene is flanked by protein-coding genes that do not show significant upregulation at competence (Figure 4A, B). Multiple OCRs flanking and within the *CLOCK* locus are enriched for TFBMs of highly expressed TF families, including Sp/Klf and other zinc-finger, bHLH and Antp (Supplementary file 15). Notably, the two closest flanking OCRs become less accessible following settlement, coincident with *CLOCK* downregulation, suggesting that they function as activating *cis*-regulatory elements. CLOCK, together with many TFs predicted to bind these regulatory regions, is enriched in larval epithelial cell types that express light-detecting cryptochromes (Rivera et al., 2012) (Supplementary file 7), consistent with a role in responding to diminishing light at sunset and being associated with the acquisition of competence.

**Figure 4.**
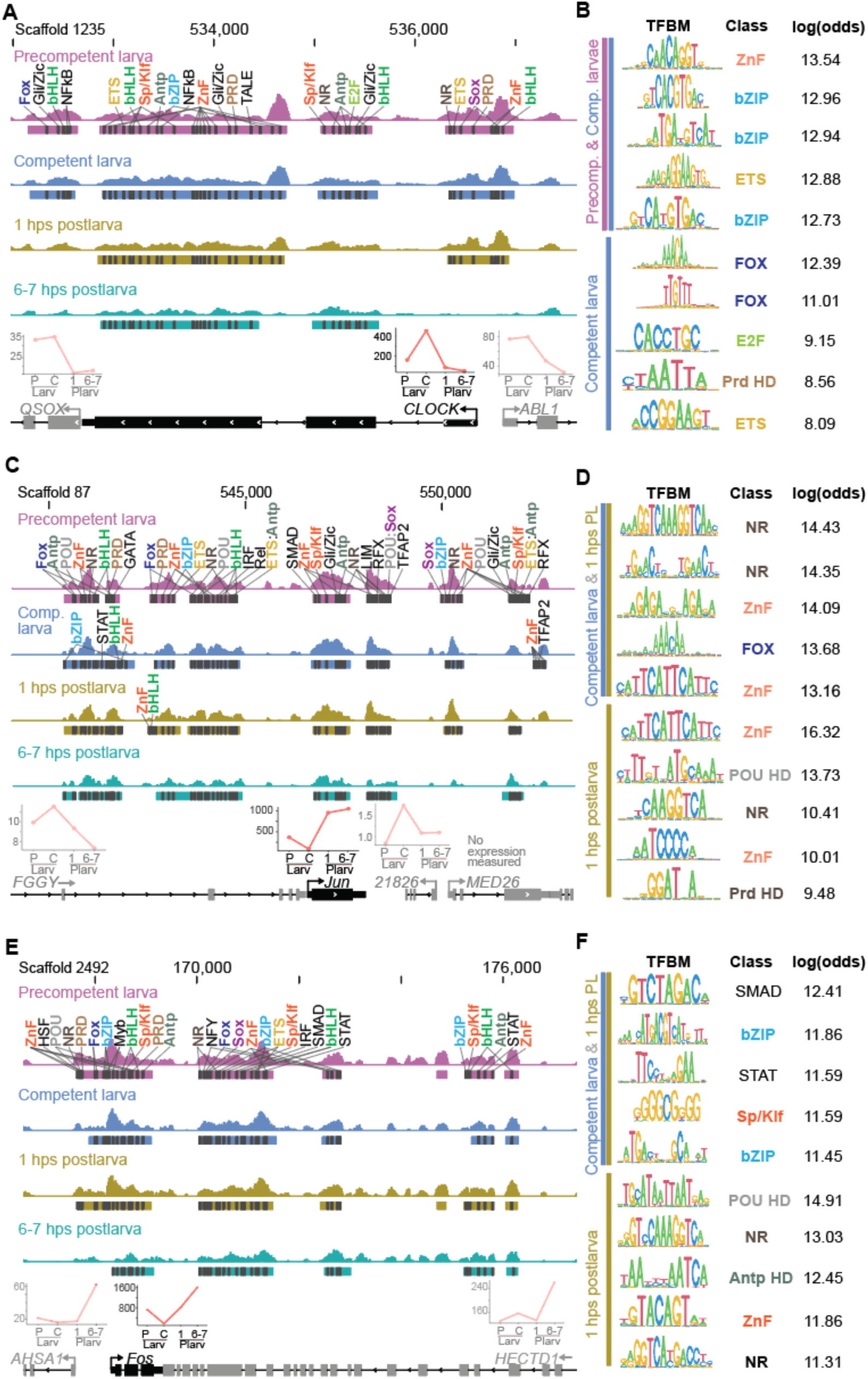
TFBMs enriched in dynamic OCRs and in *CLOCK*, *Jun* and *Fos*. A, TFBM families mapped to *CLOCK* proximal OCRs (coloured bars under ATAC-seq peaks) in larval and postlarval stages (Supplementary file 15). The locations of the putative TFBMs are demarcated by a black bar and annotated at the stage at which the OCR first appears. TFBMs colour-coded in bold correspond to highly expressed TF families. Log (odds) scores are positively correlated with motif match confidence. Protein-coding gene models at the bottom. Arrows, TSSs; chevrons, direction of transcription; exons, thick lines; UTRs, intermediate lines; introns, thin lines. The expression profiles are above the three genes (DESeq2 normalised counts). **B,** The TFBM families in *CLOCK* OCRs with the highest matches with defined motifs (higher log-odds ratio being more likely a functional binding site). **C,** TFMB families mapped to *Jun* proximal OCRs (Supplementary file 16). See description in **A** for details. **D,** The TFBM families in *Jun* OCRs with the highest matches. See description in **B** for details. **E, F,** TFBM families mapped to *Fos* proximal OCRs (Supplementary file 17). See descriptions in **A, B** for details.

AP-1 components Jun and Fos, together with other potential bZIP partners (CEBP, CREB, PAR, MAF and NFE2) (Jindrich and Degnan, 2016; Reinke et al., 2013; Rodríguez-Martínez et al., 2017), are significantly and transiently upregulated during the first hour of metamorphosis. At 1 and 6–7 hps, 470 and 340 differentially expressed genes, respectively, have AP-1 or related bZIP TFBMs within associated OCRs, including numerous other TFs and conserved developmental regulators (Supplementary file 14). Given this and that (i) Jun and Fos play a conserved role in early immediate transcriptional responses to diverse endogenous and exogenous signals, often via MAPK signalling pathway, and (ii) we have shown previously that MAPK is necessary for *A. queenslandica* settlement and metamorphosis (Bejjani et al., 2019; Eferl and Wagner, 2003; Shaulian and Karin, 2002; Ueda et al., 2016), we focussed on the OCRs in the vicinity of these two genes.

*Jun* and *Fos* are both flanked by genes with distinct expression dynamics (Figure 4C-F). OCRs associated with both loci are enriched for binding motifs of TFs that are highly expressed during competence and early metamorphosis (Supplementary files 16 and 17). These include multiple bHLH and bZIP motifs, consistent with coordinated activity of bHLH and bZIP family transcription factors during the induction of *Jun* and *Fos* at the onset of metamorphosis, and with Jun, Fos and other bZIP factors contributing to their subsequent downregulation at competence and again at 6– 7 hps. With the exception of one flanking OCR at each locus that becomes inaccessible at 6–7 hps, chromatin accessibility in the vicinity of *Jun* and *Fos* remains largely stable during larval development and early postlarval stages, suggesting that transcriptional dynamics at these loci are primarily driven by changes in TF availability rather than wholesale chromatin remodelling.

### CLOCK and other competence-associated transcription factors are repressed when larvae are prevented from experiencing sunset

To test the role of diminishing light at sunset in regulating gene expression during the acquisition of larval competence, we exposed larvae to constant light beyond sunset, a treatment that inhibits settlement and metamorphosis (Say and Degnan, 2020). Relative to larvae that experienced natural diminishing light, constant light-exposed larvae differentially up- and downregulated 779 and 879 protein-coding genes, respectively (DESeq2; *p*-adj < 0.1), activating metabolic pathways that are not normally expressed in swimming larvae (Figure 5A, B; Supplementary file 18). This transcriptional profile is distinct from both precompetent and competent larval states. Importantly, we have previously shown that larvae exposed to constant light beyond sunset swim normally, and will rapidly settle and initiate metamorphosis as normal once they are transferred into darkness (Say and Degnan, 2020), suggesting that the constant light phenotype is not due to abnormal physiology or stress; no stress pathways are enriched in larvae exposed to constant light.

**Figure 5.**
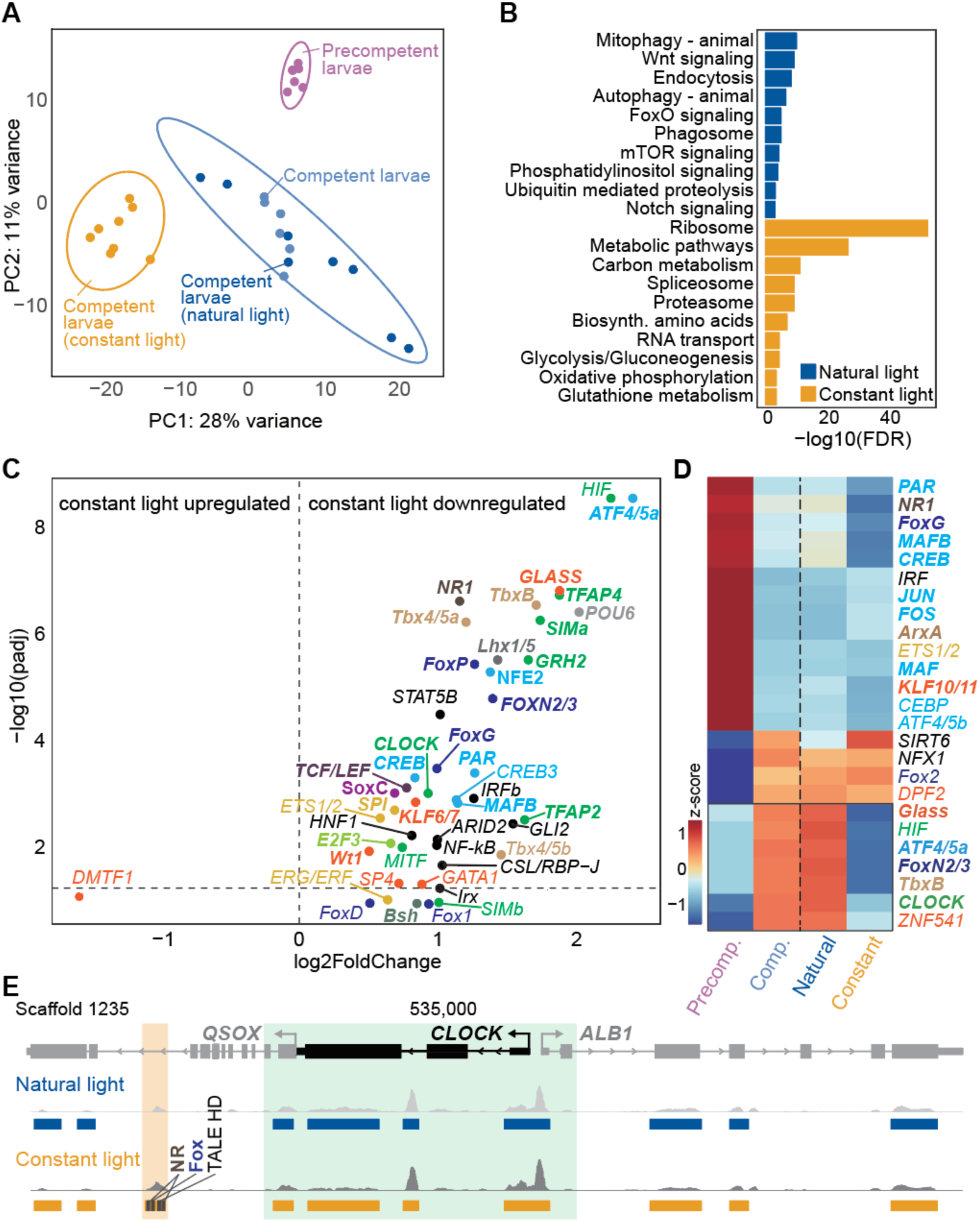
The effect of constant light on larval gene expression and chromatin state. A, PCA showing the relationship of normal transcriptomes of precompetent and competent larvae (Figure 1D), and larvae exposed to natural and constant light. **B,** Ten top KEGG categories in larvae exposed to natural and constant light (Supplementary file 18). **C,** Volcano plot of TFs up- and downregulated in larvae exposed to constant light; bolded TFs are expressed in the top 5% of all genes. **D,** Scaled heatmap of 25 TF genes that are normally differentially expressed between precompetent and competent larvae, compared to their expression in natural and constant light. TFs that are normally markedly upregulated in competent larvae but repressed by constant light are boxed (bottom). The TFs that are within the top 5% are in bold. **E,** Proximal *CLOCK* OCRs do not change in light-exposed larvae (green box; see Figure 4A), but a downstream chromatin region in a QSOX intron becomes more accessible in larvae exposed to constant light (tan box). TFBMs present in this OCR are shown as per Figure 4.

Strikingly, TF expression was broadly repressed under constant light: 45 TFs were significantly downregulated and only a single TF was upregulated (Figure 5C). Consistent with this widespread decrease in TF expression, 88% of the OCRs present in larvae under natural light conditions became inaccessible in constant light-exposed larvae (Supplementary file 18). Together, these results indicate that even brief exposure of larvae to constant light beyond sunset is associated with a novel transcriptional and chromatin state that is distinct from normal developmental trajectories, and extinguishes their capacity to respond to the algal cue, thereby preventing settlement and metamorphosis.

Several transcription factors that are normally upregulated and highly expressed in competent larvae, including CLOCK, HIF, TbxB, FoxN2/3, ATF4/5 and Glass (with CLOCK, TbxB, FoxN2/3, ATF4/5 and Glass ranking in the top 5% of expressed genes, and HIF in the top quartile), were all significantly downregulated in constant-light–exposed larvae (Figure 5D). In addition to HIF, the bHLH–PAS transcription factor SIMa, a potential partner of CLOCK, is also highly expressed in competent larvae and significantly repressed under constant light. Consistent with CLOCK and its potential partners having pioneer-like regulatory properties related to metamorphosis, a large proportion (24%) of the OCRs that close in constant light-exposed larvae have bHLH-PAS and E-box family motifs.

Examination of chromatin accessibility across the CLOCK locus and within 100 kb of its genomic neighbourhood revealed that the proximal OCRs normally associated with *CLOCK* at competence (Figure 4A) remain largely unchanged under constant light, whereas six distal regulatory regions exhibit altered accessibility, with five becoming less accessible (Supplementary file 18). Notably, the sole region that increases accessibility in constant-light larvae lies within an intron of an adjacent gene approximately 2.5 kb downstream of *CLOCK* (Figure 5E). These rapid, light-dependent changes in chromatin state support the existence of a conditional gene regulatory program that is normally suppressed at sunset. When activated by aberrant light conditions this program downregulates *CLOCK* and other competence-associated TF genes, disrupting chromatin priming, and preventing the transcriptional and chromatin changes required for larval competence, settlement and metamorphosis (Figure 6).

**Figure 6.**
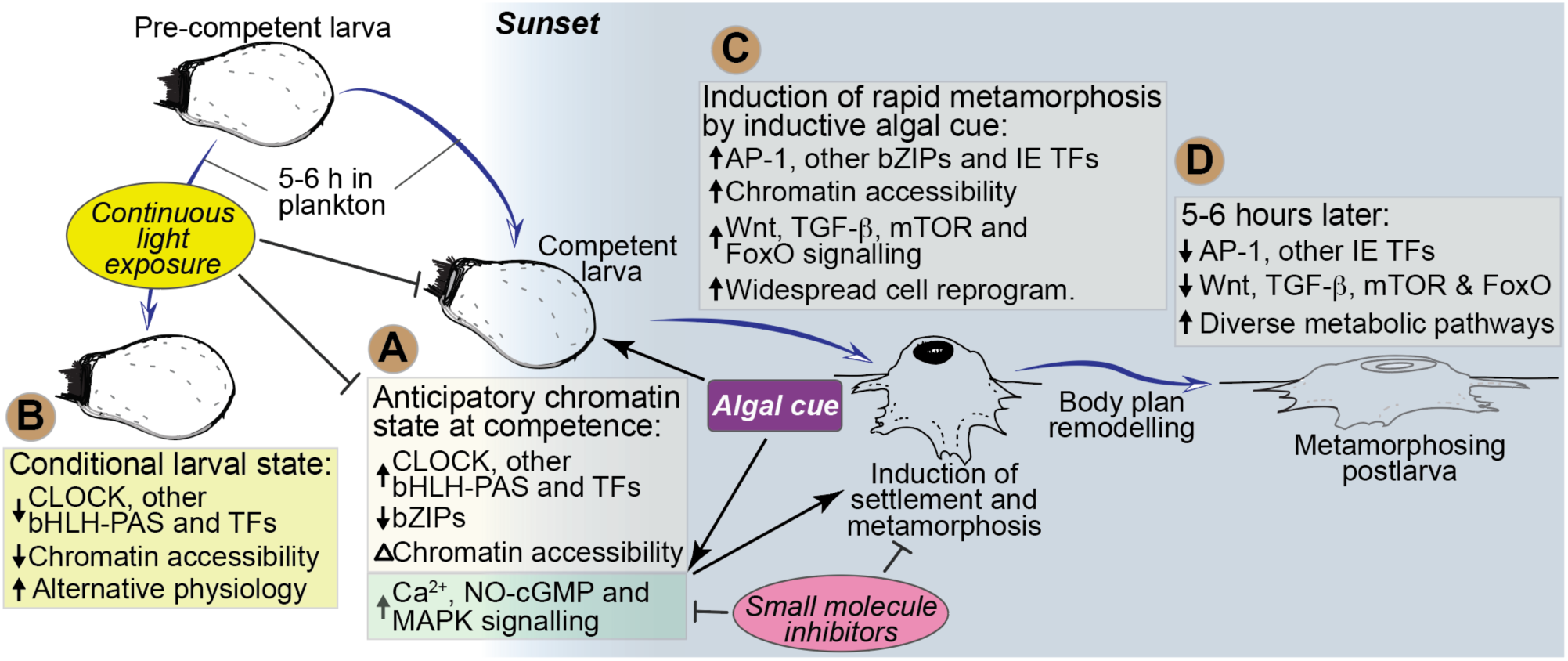
Environmental cues, genomic regulatory processes and signalling events underlying larval competence and early metamorphosis in *A. queenslandica*. This model combines results from this and previous studies (Nakanishi et al., 2015; Say and Degnan, 2020; Ueda et al., 2016). **A,** Pre-competent larvae emerge from adults early in the afternoon, then swim in the water column for several hours before becoming competent to settle and initiate metamorphosis immediately after sunset (Degnan and Degnan, 2010; Say and Degnan, 2020). With the acquisition of competence is: (i) the upregulation of genes encoding CLOCK and other bHLH–PAS TFs, and other TFs, and the downregulation of AP-1 and other bZIPs; and (ii) a change in chromatin accessability in the vicinity of genes that are differentially expressed at competence and in the first hour of metamorphosis. **B,** Larvae prevented from experiencing sunset do not settle in the presense of inductive algae nor initiate metamorphosis (Say and Degnan, 2020). Chromatin accessibility in light-exposed larvae decreases markedly and larvae move into a alternative state that allow for prolonged periods in the plankton. This includes the repression of over 40 TF genes, including *CLOCK* and *HIF*. **C,** A cue on the surface of the alga *A. fragilissima* induces an internal signalling cascade that includes activation of calcium (Ca^2+^), nitric oxide (NO-cGMP) and MAPK pathways; larvae exposed to pharmacological agents that inhibit these pathways do not settle and metamorphose (Nakanishi et al., 2015; Song et al., 2020; Ueda et al., 2016). Newly settled postlarvae undergo large changes in chromatin accessability and gene expression within the first hour of settling, repressing larval genes and activating multiple developmental signallng pathways and over 60 TFs, including AP-1 and other potential partner bZIP TFs, and other conserved Immediate-early (IE) TFs. During this first hour there are rapid and extensive morphogenetic changes with multiple cell types being reprogrammed (Blard et al., 2026). **D,** 5-6 hour later, most developmental TF and signalling genes are downregulated, and diverse metabolic genes are activated, indicative of an overall shift in postlarval developmental and physiological state.

## Discussion

Metamorphosis represents one of the most dramatic and consequential transitions in an animal’s life, transforming planktonic larvae specialized for dispersal and environmental sensing into benthic juveniles/adults adapted for growth, survival and reproduction. In *Amphimedon queenslandica*, this transition is both environmentally induced and remarkably rapid, unfolding within hours of settlement. We show that this swift morphological transition is accompanied by an unexpectly rapid and large transcriptional response in the first hour of metamorphosis, with the expression level of over 4850 genes changing significantly. Such speed implies that genomic regulatory processes must be initiated before the inductive cue is encountered. Here, by integrating time-resolved transcriptomic and chromatin accessibility profiling across larval competence, settlement, and early postlarval development, we show that larvae undergo extensive regulatory reorganisation while still swimming in the plankton, prior to metamorphic induction. In particular, diminishing light at sunset is associated with widespread changes in chromatin accessibility that precede large-scale transcriptional activation at settlement (Figure 6). These findings indicate that the larval genome enters an anticipatory state that is permissive for rapid, coordinated transcriptional reprogramming, rather than responding de novo to the inductive environmental cue (Nakanishi et al., 2015; Say and Degnan, 2020; Ueda et al., 2016).

A defining feature of *A. queenslandica* metamorphosis is the immediate and substantial activation of a large suite of deeply conserved developmental transcription factors (TFs). To our knowledge, such a diverse suite of TFs reaching expression levels within the top 5% of all genes has never before been observed in animal developmental transitions. Their exceptionally high expression at competence and at the onset of metamorphosis may confer regulatory robustness, ensuring sustained occupancy of *cis*-regulatory elements and enabling rapid, coordinated shifts in gene expression. Among these, the AP-1 components Fos and Jun, together with their potential bZIP heterodimeric partners (Reinke et al., 2013; Rodríguez-Martínez et al., 2017), are rapidly and strongly induced following settlement. AP-1 is a conserved effector of environmentally responsive signalling pathways, frequently activated downstream of MAPK cascades to regulate cell differentiation, proliferation and death (Bejjani et al., 2019; Eferl and Wagner, 2003; Shaulian and Karin, 2002). The strong enrichment of bZIP motifs within open chromatin regions associated with genes that change expression immediately after settlement supports a central role for AP-1 in coordinating the transcriptional reprogramming underlying metamorphosis. Consistent with this interpretation, inhibition of MEK/ERK signalling blocks settlement and metamorphosis in *A. queenslandica* and other marine invertebrate larvae (Cavalcanti et al., 2020; Heyland and Moroz, 2006; Shikuma et al., 2016; Taylor and Heyland, 2018; Ueda et al., 2016). Thus, although the timing of competence acquisition and the identity of inductive settlement cues are largely species specific, the initiation of metamorphosis itself may rely on a conserved regulatory logic involving MAPK signalling and AP-1–mediated transcriptional activation.

The induction of metamorphosis in *A. queenslandica* is conditional on the prior acquisition of competence, which develops in larvae after a period in the planktonic and exposure to darkness at sunset (Degnan and Degnan, 2010; Say and Degnan, 2020). We show here that competence acquisition is accompanied by extensive remodelling of chromatin accessibility and changes in gene expression. These changes not only appear to permit settlement in response to the algal cue, but also establish a permissive chromatin landscape that enables the rapid and large-scale transcriptional response observed during the first hour of metamorphosis (Figure 6). Larvae prevented from experiencing sunset fail to settle and metamorphose, exhibit markedly reduced expression of CLOCK and multiple other highly expressed developmental TFs, and display widespread loss of chromatin accessibility. Together, these findings support a model in which CLOCK and associated pioneer-like TFs mediate anticipatory chromatin priming that is required for the subsequent induction of AP-1 and other developmental regulators at settlement.

Diminishing light intensity has been implicated in regulating larval settlement in a range of marine species (Thorson, 1964), raising the possibility that circadian entrainment of competence via CLOCK represents a widespread and ancestral feature of the pelagobenthic life cycle. Notably, *A. queenslandica* larvae maintained in constant light do not simply remain in a pre-competent state but instead transition into an alternative transcriptional and chromatin configuration that is distinct from both pre-competent and competent larvae. This observation suggests the existence of sensitive, rapidly deployable and conditional gene regulatory programs that are normally suppressed under predictable sunset conditions. Activation of such alternative programs may allow larvae to delay settlement and extend dispersal when environmental conditions are unfavourable, thereby modulating connectivity and colonisation potential.

Together, our results support a model in which environmentally predictable planetary cues, such as the daily light–dark cycle, are integrated during the planktonic larval phase to establish a permissive genomic state that enables rapid induction of metamorphosis upon encountering a suitable, often species-specific, settlement cue. In *A. queenslandica*, this state is characterised by (i) extensive chromatin remodeling prior to settlement, (ii) transient but exceptionally high expression of conserved developmental TFs at both competence and the initiation of metamorphosis, and (iii) enrichment of binding motifs of these TF families in regulatory regions associated with genes regulated during early metamorphosis. Preventing larvae from experiencing sunset does not simply delay development, but instead diverts them into an alternative transcriptional and chromatin configuration that is incompatible with settlement and metamorphosis, consistent with the activation of a unique conditional regulatory program. Although functional dissection of individual regulators is currently unfeasible in *A. queenslandica*, the temporal ordering of chromatin accessibility changes, TF expression, and environmental induction suggests light-entrained transcriptional activity—potentially involving CLOCK-associated and AP-1/bZIP family factors—contributes to chromatin priming and rapid transcriptional reprogramming. We propose that such anticipatory chromatin regulation may represent a general solution to the problem of speed and robustness in metamorphosis, enabling biphasic animal life cycles to integrate predictable planetary signals with species-specific ecological cues across evolutionary timescales.

The existence of an alternative, inducible regulatory program that is conditional on the environmental experience of the sponge larva provides a potential mechanistic explanation for how larvae modulate dispersal range and timing of settlement. The magnitude and tempo of these regulatory changes are comparable to the rapid metamorphic transitions observed across many marine invertebrates (Hadfield, 2000; Hadfield et al., 2001), suggesting that the genomic control mechanisms identified here may be broadly relevant. However, in the absence of equivalent chromatin-resolved datasets from parahoxozoan lineages (e.g. echinoderms, hemichordates, annelids, molluscs, bryozoans or cnidarians), the extent to which these mechanisms are conserved remains an open question. We propose that, despite the extraordinary diversity of larval forms and settlement cues across the animal kingdom, metamorphosis is governed by a shared set of genomic control principles that integrate planetary and species-specific environmental signals through deeply conserved transcriptional regulators.

## Methods

### Larval and postlarval collection and cell isolation

Adult *Amphimedon queenslandica* were collected from the intertidal reef flat of Shark Bay, Heron Island Reef, Southern Great Barrier Reef (23° 27’S, 151° 55’E) and were immediately transferred to aquaria at Heron Island Research Station, where they were maintained in flow-through, unfiltered ambient seawater drawn from the adjacent reef flat (Leys et al., 2008). Naturally emerging larvae were collected in the early to mid-afternoon, and maintained in ambient conditions as previously described (Say and Degnan, 2020). Randomly-selected larvae were transferred at sunset (5-6 hours post-emergence; hpe) to a 6-well dish with 10 ml of 0.22 µm filtered seawater (FSW) (10 per well), and then exposed 30 min later to three small branches of the coralline algae *Amphiroa fragilissima* as previously described (Say and Degnan, 2020). These dishes were maintained under ambient water temperature and light conditions (Say and Degnan, 2020). For collection of postlarval stages, dishes were checked every 30 mins to determine the time of settlement, defined by the larva remaining attached to the alga when the dish was given a gentle shake.

Six randomly-sampled larvae and postlarvae were collected from each of the following stages for cell dissociation, and CEL-seq2 and ATAC-seq analyses: precompetent larvae (0-1 hpe); competent larvae (5-6 hpe); 1 hps postlarvae; 3-4 hps postlarvae; 6-7 hps postlarvae; and 11-12 hps postlarvae (Figure 1D). All individuals were treated individually (i.e. not pooled). All postlarvae that had settled on *A. fragilissima* were removed rapidly from the alga with a sharp tungsten needle and transferred into FSW on ice. Precompetent and competent larvae were treated the same but did not need to be removed from the alga. All individual larvae and postlarvae were washed three times for 2 min in ice cold FSW and then centrifuged through a 25 µm metal mesh fitted into a 1.5 ml Eppendorf at 500 rpm for 10 min at 4°C. Pelleted fragments and cells were resuspended and washed with ice cold 0.22 µm filtered calcium-magnesium-free artificial seawater (CMFSW; 449 mM NaCl, 33 mM Na2SO4, 2.15 mM NaHCO3, 9 mM KCl, 10 mM Tris-HCl pH 10, and 2.5 mM EGTA) for 5 min and recentrifuged at 500 rpm for 10 min at 4°C. The cell pellet was resuspended in 20 µl ice cold CMFSW. This procedure took less than 25 minutes at 4°C, thus minimising changes in gene expression or chromatin state associated with these perturbations.

For four developmental stages – precompetent and competent larvae, and 1 and 6-7 hps postlarvae – we separately assayed both gene expression and chromatin accessibility in the same six individuals by splitting their resuspended cells into two pools. For each individual of these four stages, 5 and 15 µl of the suspended cells were used for CEL-seq2 and ATAC-seq library construction, respectively. For CEL-seq2, 5 µl of the cell suspension was immediately mixed with 50 μl ice-cold TRIZol and stored in -80°C. For ATAC-seq, 15 µl of the cell suspension was immediately mixed with 50 μl ice cold ATAC-seq cell lysis buffer and processed as described below. The other two stages – 3-4 and 11-12 hps postlarvae – were assayed only for gene expression, so that total 20 µl of the cell suspension was immediately mixed with 200 μl ice-cold TRIZol, resulting in the same dilution as above, and stored in -80°C.

### Imaging early metamorphosis

Images of *A. queenslandica* postlarvae were taken with a Nikon SMZ25 microscope using the program Nikon Elements and 6X zoom every 3 min, starting from ∼10 min after the initiation of settlement on *A. fragilissima* through to ∼ 2 hps. Each time point consisted of a z-stack with 14 images at an interval of 25 µm, which were then stacked into a single image using the focus stacking program Zerene Stacker with the P Max algorithm (Brecko et al., 2014). These were cropped, brightened and denoised, then unsharp masked using Adobe Photoshop. To generate the timelapse movie, the frames were compiled into an avi using ImageJ and converted to mp4 (or mkv) using AnyVideoConverter (https://www.any-video-converter.com/en8/for_video_free/).

### CEL-seq2 library construction, sequencing and analysis

RNA isolation and quality-assessment, and CEL-seq2 library construction all were performed as previously described (Hashimshony et al., 2016, 2012; Sogabe et al., 2019). Libraries were sequenced on two lanes of Illumina HiSeq-PE 150 platform and raw sequencing data were processed using a publicly available pipeline (https://github.com/yanailab/celseq2) (Hashimshony et al., 2016) with a built-in a suite of python-based scripts for demultiplexing, mapping to the genome, and counting reads. Raw reads were mapped to *A. queenslandica* Aqu3.1 genome (NCBI Bioproject PRJNA668660) (Xiang et al., 2022) using Bowtie2 (Langmead and Salzberg, 2012), and assessed for coverage and quality as previously described (Say and Degnan, 2020; Sogabe et al., 2019) (Supplementary file 1). Count tables for the two sequencing lanes were merged using a custom R script (https://github.com/hfyuanuq/thesis.scripts/blob/main/CEL-Seq2- combine%202%20lanes) before the differential expression analysis. CEL-seq2 raw counts from the two larval ages in two different light regimes were obtained from https://github.com/tahshasay/TS1015_DESeq2/tree/master (Say and Degnan, 2020) and larval single-cell RNA-seq raw counts were obtained from https://github.com/tanaylab/ (Sebé-Pedrós et al., 2018a).

### Analysis of differentially expressed genes

Only genes with at least 4 read counts in at least 4 samples were considered expressed; those not meeting these criteria were filtered out from all subsequent analysis. PCA of all expressed genes was performed on variance-stabilizing transformation (VST) transformed counts and visualized using the function ’plotPCA’ to examine sample-to-sample distances and identify potential outliers with 95% confidence ellipses. Differential gene expression was conducted in R (R, 2021) using the Bioconductor package DESeq2 (Love et al., 2014a). Pairwise comparisons were conducted between each of the six developmental stages to produce a differentially expressed gene (DEG) list for each pair of stages, and an extra comparison was conducted between 1 hps and 6-7 hps, with a False Discovery Rate threshold (FDR) < 0.1 (Supplementary file 3). This FDR allowed us to detect substantial changes in transcript abundance within one hour (that is, between competent larvae and newly settled 1 hps postlarvae), taking into account known eukaryotic rates of RNA transcription and degradation (Sun et al., 2012). Venn diagrams were generated using the R package ggVennDiagram (Gao et al., 2021). Heatmaps were produced in R using packages pheatmap (https://CRAN.R-project.org/package=pheatmap) and RColorBrewer (https://CRAN.R-project.org/package=RColorBrewer) to visualize expression profiles with rlog normalised counts for CEL-seq2 datasets. Single-cell RNA-seq reads were normalised with the function CPM (*cpm(data, log=TRUE)*) using the Bioconductor package edgeR (version 3.14.0) implemented in R (Robinson et al., 2010). Alluvial (Sankey) diagrams were generated using SankeyMATIC (vBeta https://sankeymatic.com/build/).

### Co-expression analysis

Weighted Gene Co-expression Network Analysis (WGCNA; version 1.61)(Langfelder and Horvath, 2008) was performed on the VST-transformed developmental dataset using R (version 4.0) (Love et al., 2014b). We used a signed Nowick-type topological overlap metric with soft threshold of *β* = 3. Co-expression modules were defined using dynamic tree cut (height 0.15; minimum module size 30). Module eigengenes were correlated with time points. To retain the distinct expression patterns of each module, a merging distance threshold of 0.1 was applied.

### ATAC-seq library preparation, sequencing and analysis

ATAC-seq libraries were prepared as previously described (Cornejo-Páramo et al., 2022). Briefly, we added ATAC-seq cell lysis buffer (10 mM Tris-HCl, pH 7.4, 10 mM NaCl, 3 mM MgCl2, 0.1% v/v IGEPAL CA-630) to the 15 µl of dissociated cells (15-20,000 cells) and centrifuged at 500 g for 10 min at 4°C. The pellet was resuspended in 50 μl transposition reaction and incubated at 37°C for 30 min. The resultant DNAs were amplified, purified and quality assessed as previously described (Cornejo-Páramo et al., 2022). Libraries were sequenced on a 2 x 101 bp NovaSeq SP flow cell platform. Raw reads were trimmed and low-quality reads removed using FastQC and Trim-Galore (parameters: -q 30 --phred33 --stringency 4 --length 20 -e 0.1) (https://github.com/FelixKrueger/TrimGalore). Reads were mapped to the *A. queenslandica* Aqu3.1 genome using Bowtie2 (parameters: -X 1000) (Langmead and Salzberg, 2012). Paired reads were identified using SAMtools (parameter: -f 2) (Li et al., 2009) and duplicated reads were identified using Sambamba and removed (Danecek et al., 2020). MACS2 was used to call peaks (parameters: -g 1.2e8 --nomodel --shift -100 --extsize 200) (Zhang et al., 2008). This yielded over 46 million reads that corresponded to over 20,000 peaks from each library (Supplementary file 8). ATAC-seq library quality was determined by calculating fraction of reads in peaks (FRiP) scores using the ENCODE ATAC-seq pipeline (https://www.encodeproject.org/atac-seq/); a FRiP score over 0.2 was deemed acceptable (Supplementary file 8). Peaks from the three replicate ATAC-seq libraries for each stage were pooled and overlapping peaks were intersected using bedtools (Quinlan and Hall, 2010), also in the ENCODE ATAC-seq pipeline (https://www.encodeproject.org/atac-seq/).

### Analysis of open chromatin regions

ATAC-seq peaks were annotated using the Bioconductor package ChIPseeker (Yu et al., 2015), with the promoter region set as ± 500 bp (Supplementary file 9). The Bioconductor package DiffBind (V2.16.0) was used for differential binding analysis of peaks (Stark and Brown, 2011). Significantly different peaks (FDR < 0.05) between the four developmental stages were identified by pairwise comparisons on successive stages using DESeq2 (Love et al., 2014a). Consistent peaks and overlapping peaks were generated in DiffBind using the function dba.peakset (parameters: consensus=DBA_FACTOR, minOverlap=0.33) (Stark and Brown, 2011). Library correlation analysis was performed using ChromVAR (Schep et al., 2017) to determine peaks across all 12 libraries. Genes associated with these peaks were compared to the list of genes associated with ChIP-Seq peaks (Gaiti et al., 2017b).

### Motif identification, annotation, clustering, and accessibility analysis

Homer (Heinz et al., 2010) was used to identify TFBMs with parameters of -size 500 -len 6, 8, 10, 12 -h. All identified motifs were combined into a single motif list, and we excluded all the potential *A. queenslandica* specific, sequence bias motifs, and non-significant identified motifs (p-value > 1e^- 10^). Motifs were matched to the database (https://jaspar2022.genereg.net/downloads/) that we customised by removing fungi- and plant-specific motifs (available at https://github.com/hfyuanuq/atac-pub) using MEME-TomTom (version 5.5.8) with default threshold 0.5 (Gupta et al., 2007). After motif matching, all putative TFBMs were assigned with JASPAR motif identifiers. Then motifs were clustered into TF classes and families based on JASPAR annotations using the Bioconductor package TFBSTools (Tan and Lenhard, 2016).

TF binding sites analysis was performed using the GimmeMotifs scan tool (Bruse and Heeringen, 2018; Heeringen and Veenstra, 2011). OCR sequences of expressed genes and differentially expressed genes were scanned against the JASPAR non-redundant database. Motif scan significance was determined using log-odds scores calculated relative to the *A. queenslandica* genomic background model. Putative motif occurrences were filtered using a FDR threshold of 0.01. Following the scan, all putative TFBMs were clustered into TF class and family as previously described. Chromatin accessibility and peak profiles were visualized using the R package Gviz (Hahne and Ivanek, 2016) and the Integrative Genomics Viewer (IGV) software (Thorvaldsdóttir et al., 2013).

### Enrichment analyses

KEGG enrichments, which are threshold-dependent, were performed using the online tool STRING (https://string-db.org/) with default settings. To more fully capture functions that are enriched among highly regulated genes, rather than only among highly expressed genes, we also conducted a GO_MWU (Mann-Whitney U test for Gene Ontology term enrichment)analysis, based only on fold-change (Voolstra et al., 2011). Available at: https://github.com/z0on/GO_MWU).

### Light perturbation experiment

From adults collected maintained in flow-through seawater aquaria at Heron Island Research Station, as described above, > 50 naturally-released larvae were collected within 30 min of emerging, and split equally among two 500 ml glass beakers of sea water. One beaker was exposed to natural light and the other to constant light, both at ambient temperature, as described previously (Say and Degnan, 2020). At sunset (5–6 hpe), eight larvae from each beaker were randomly collected. Their cells were dissociated and CEL-seq2 and ATAC-seq libraries were constructed, sequenced and analysed as described above (Yuan et al., 2025).

CEL-seq2 and ATAC-seq sequencing were performed using the Illumina NovaSeq X Series sequencer (PE150, NovaSeq Control Software v1.2.2.48004 and Real Time Analysis v4.6.7). Precompetent and competent larval, and constant and natural light CEL-seq2 datasets were combined and integrated, with competent larval and natural light samples being combined into a single group. Batch effects were corrected using ComBat-Seq (Zhang et al., 2020). Differential OCRs and associated TFBMs were identified, characterised and visualised as described above. For each motif, the average deviation difference between constant and natural conditions was computed to determine the direction and magnitude of activity change.

## Data availability

CEL-seq2 and ATAC-seq data for normal light condition are available in the NCBI database under BioProjects PRJNA1162246 and PRJNA1161795, respectively, and for constant light condition under BioProjects PRJNA1394668 and PRJNA1394227, respectively.

## Supporting information

Suppfile1

Suppfile2

Suppfile3

Suppfile4

Suppfile5

Suppfile6

Suppfile7

Suppfile8

Suppfile9

Suppfile10

Suppfile11

Suppfile12

Suppfile13

Suppfile14

Suppfile15

Suppfile16

Suppfile17

Suppfile18

## Acknowledgements

This study was supported by funds from the Australian Research Council to S.M.D and B.M.D (DP210100703 and DP230102109). Biological samples were collected with the support of the Heron Island Research Station and Bin Yang, and ATAC-seq was performed under guidance from Miloš Tanurdžić. Computational analyses were enabled by support from the Queensland Cyber Infrastructure Foundation, with a special thanks to Nick Rhodes.

## Author contributions

S.M.D, B.M.D. and H.Y. conceived and designed the project. H.Y. isolated cells and prepared all libraries. H.Y. performed all gene expression and chromatin state analyses. O.B. and Z.P. visually documented postlarval morphogenesis. H.Y, S.M.D. and B.M.D wrote the manuscript with minor contributions from the other authors.

## Materials & Correspondence

Materials & Correspondence should be addressed to S.M.D or B.M.D.

---

**Figure 1 – figure supplement 1.** Larval and postlarval gene expression. A, WGCNA co-expression modules of genes expressed through larval and postlarval development. Number of coding genes and TFs are shown. **B, C,** KEGG (B) and GO-MWU (C) analysis of WGCNA co-expression modules that comprise genes that are down (blue, pink) and up (red, yellow-green, tan) regulated at the start of metamorphosis. BP, biological process; CC, cellular component; and MF, molecular function. **D,** Developmentally enriched GO-MVU terms (-log p-value; maximum of 10 shown) based on significantly upregulated genes at each developmental stage. **E, F,** Scaled heatmaps of expression of TFs in larval cell types previously defined using scRNA-seq (Sebé-Pedrós et al., 2018). **E,** TF genes significantly differentially expressed between precompetent and competent larval stages. **F,** TF genes significantly differentially expressed between competent larval and 1 hps postlarval stages. Highly expressed TFs are in bold, and arrows in E and F show whether the TF is up or downregulated between successive stages.

**Figure 2 – figure supplement 1.** ATAC-seq analysis of *A. queenslandica* larval and early postlarval chromatin. Analyses were performed on cells dissociated from three individual precompetent and competent larvae, and 1 and 6-7 hps postlarvae. **A,** Insert size distribution of ATAC-seq libraries. Three biological replicates for each stage shown. **B,** Bar graph of number of ATAC-seq peaks in each replicate. Developmental stages are colour-coded as per Fig. 1. **C,** Bar graph of the total number of OCRs identified per developmental stage. **D,** Bar graph of the number of ATAC-seq peaks shared between larval and postlarval stages. Most peaks are present in all four stages. **E,** Bar graph of the number of ATAC-seq peaks shared between stages. **F,** ATAC-seq FRiP scores for all replicates. Dotted lines, ENCODE standards (https://www.encodeproject.org/atac-seq/). **G,** Boxplot of ATAC-seq peak insert size, calculated using the arithmetic mean across replicates for each stage. **H,** Boxplot of ATAC-seq peak width size, calculated as per h. **I,** Heatmaps of total ATAC-seq reads for each developmental stage within 1 kb up and downstream of TSSs of all genes. **J,** Normalised ATAC-seq read density from TSS to transcription end site (TES) and 5 kb up and downstream for all genes in each replicate and stage, showing enrichment and depletion at TSSs and TESs, respectively. **K,** Chromatin accessibility read density 1 kb up and downstream of the TSS, by stage. Top panel plots the read density for three replicate ATAC-seq analyses for each larval and postlarval stage. Bottom panel shows the consensus plot for each stage. **L,** The percentage of ATAC-seq peaks mapping to genomic features for larval and postlarval replicates. Promoter region is defined as a region within 500 bp of the TSS. Downstream is defined as ≤300 bp of the end of a gene. **M,** The percentage of ATAC-seq peaks up and downstream of the TSS for larval and postlarval replicates, partitioned into classes based on distance from the TSS.

**Figure 3 – figure supplement 1.** Chromatin dynamics through larval and early postlarval development. A, PCA of the 3,482 DACRs that significantly change between in least one larval or postlarval stage (DESeq2; *p*-adj < 0.1); n = 3 for all stages. **B,** Hierarchical clustered heatmap of Pearson correlation of replicated larval and postlarval stages based on log10-transformed read counts of the 3,482 DACRs. **C,** Examples of the location and frequency of ATAC-seq reads mapped to the genome are shown and the regions of significant change are highlighted in grey. Scaffold locations of OCRs are shown above and gene names below. Direction of transcription is depicted by chevrons and exons are depicted by black blocks. Below gene name is stage of development in which the gene is up or downregulated, and the p-adjusted significance values of the OCRs and expression levels. *PHAF1, Phagosome Assembly Factor 1*; *ACBD5, Acyl-CoA binding domain-containing 5*; *UVSSA, UV stimulated scaffold protein A*; *WASHC2C, WASH complex subunit 2C*; *PTPRK, Protein tyrosine phosphatase receptor type K*. **D,** Venn diagram of the OCRs unique to and shared between precompetent and competent larvae, and 1 and 6-7 hps postlarvae (Supplementary fie 10). **E,** Venn diagram of the conserved TFBMs unique to and shared between precompetent and competent larvae, and 1 and 6-7 hps postlarvae as determined by Homer (Supplementary file 10). **F-H,** Analysis of TFBMs present in DACRs associated with the five WGCNA modules whose genes change across the intitiation of metamorphosis (Fig. 1I and Supplementary file 11); module colours shown. **F,** Upset plot of TFBMs unique to and shared between WGCNA modules. **G,** Heatmap of the most common TFBMs shared across all modules. TFBMs that potentially interact with highly expressed TFs is colour-coded by TF class as per Fig. 1K; black, TFBMs of other expressed TFs. Jasper code follows TF class/family name to the right. **H,** Top, the two TFBMs that are significantly enriched in OCRs of all five WGCNA modules, with Homer p-values to the right. Bottom, three TFBMs enriched in OCRS associated with genes comprising blue and pink modules.

## Supplementary files

**Supplementary file1.** Information on CEL-seq2 libraries.

**Supplementary file 2.** Normalised and average CEL-seq2 counts for all expressed genes using DESeq2.

**Supplementary file 3.** Genes that are differentially expressed during larval development and early metamorphosis.

**Supplementary file 4.** KEGG enrichments for genes that are upregulated at each larval and early postlarval stage, and MVU enrichments of differentially expressed genes between successive stages.

**Supplementary file 5.** KEGG enrichments for genes in each WGCNA modules.

**Supplementary file 6.** TF genes that are expressed and differentially expressed during larval development and early metamorphosis.

**Supplementary file 7.** Raw and normalised MARS-seq counts (Sebé-Pedrós et al., 2018) of differentially expressed TF genes in larval cell types.

**Supplementary file 8.** Information on ATAC-seq libraries: sequencing reads, mapping ratios, peaks, and fractions of reads in called peak regions.

**Supplementary file 9.** Genomic location and TFBMs enrichment in OCRs present during larval development and early metamorphosis. Larval enhancers and promoters inferred by ChIP-seq (Gaiti et al., 2017) overlapped with OCRs identified in precompetent larvae in the present study.

**Supplementary file 10.** OCRs and their TFBMs associated with expressed genes at all four stages.

**Supplementary file 11.** OCRs and their TFBMs associated with WGCNA modules.

**Supplementary file 12.** OCRs and their TFBMs associated with DEGs between precompetent and competent, and competent and 1 hps.

**Supplementary file 13.** OCRs and their PAS related (bHLH) TFBMs associated with differentially expressed genes during larval development and early metamorphosis.

**Supplementary file 14.** OCRs and their Fos/Jun-related (bZIP) TFBMs associated with differentially expressed genes during larval development and early metamorphosis.

**Supplementary file 15.** OCRs and TFBMs found in and around *CLOCK* during larval development and early metamorphosis.

**Supplementary file 16.** OCRs and TFBMs found in and around *Jun* during larval development and early metamorphosis.

**Supplementary file 17.** OCRs and TFBMs found in and around *Fos* during larval development and early metamorphosis.

**Supplementary file 18.** Comparison of gene expression, chromatin state and accessible TFBMs in larvae exposed to natural and constant light.

