## Supplementary figures and images for "Light-entrained chromatin priming poises rapid metamorphosis in a marine sponge"

### Fig 1 supp 1

#### Supplementary Figure 1

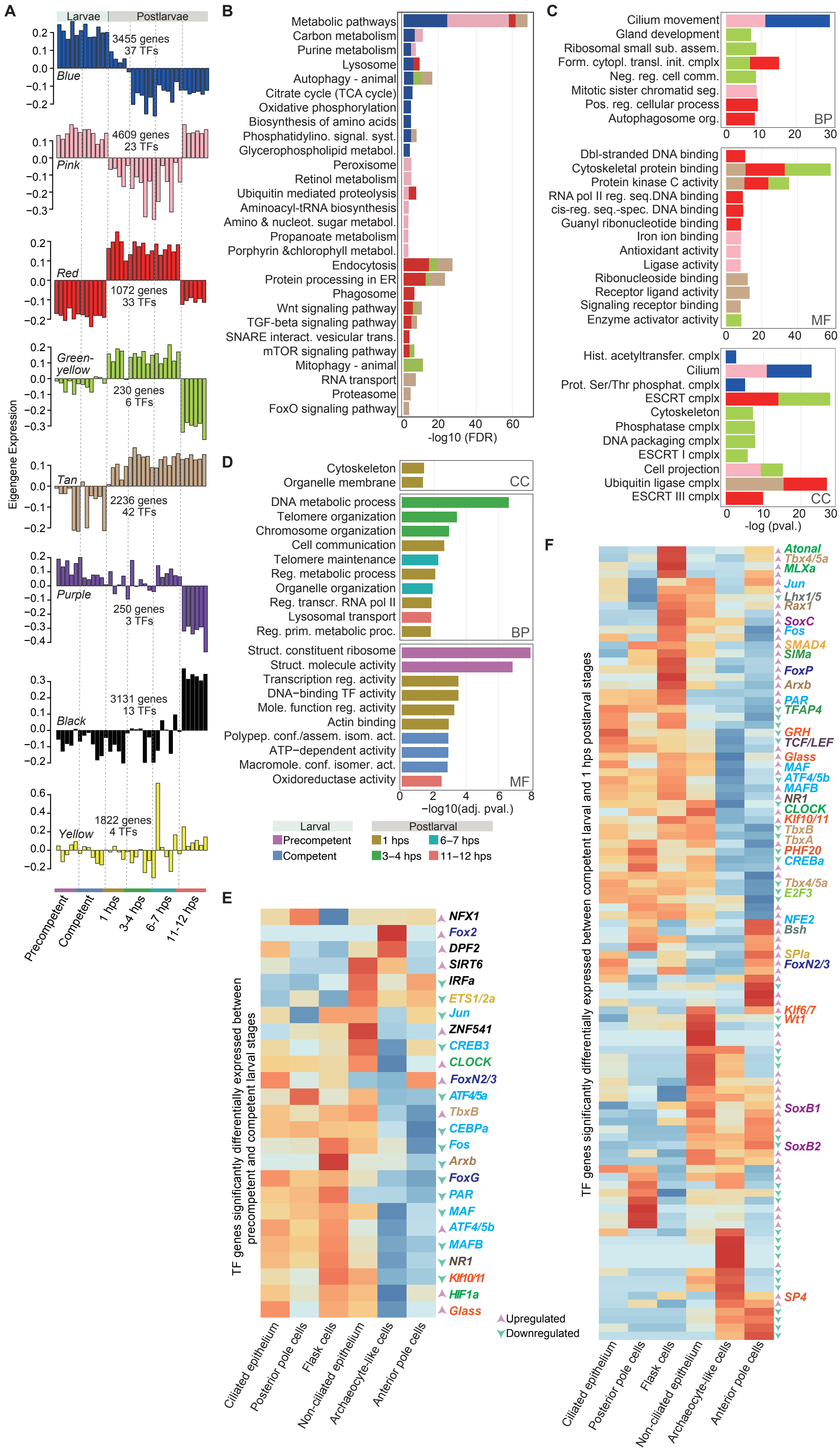

### Fig 2 supp 1

#### Supplementary Figure 2

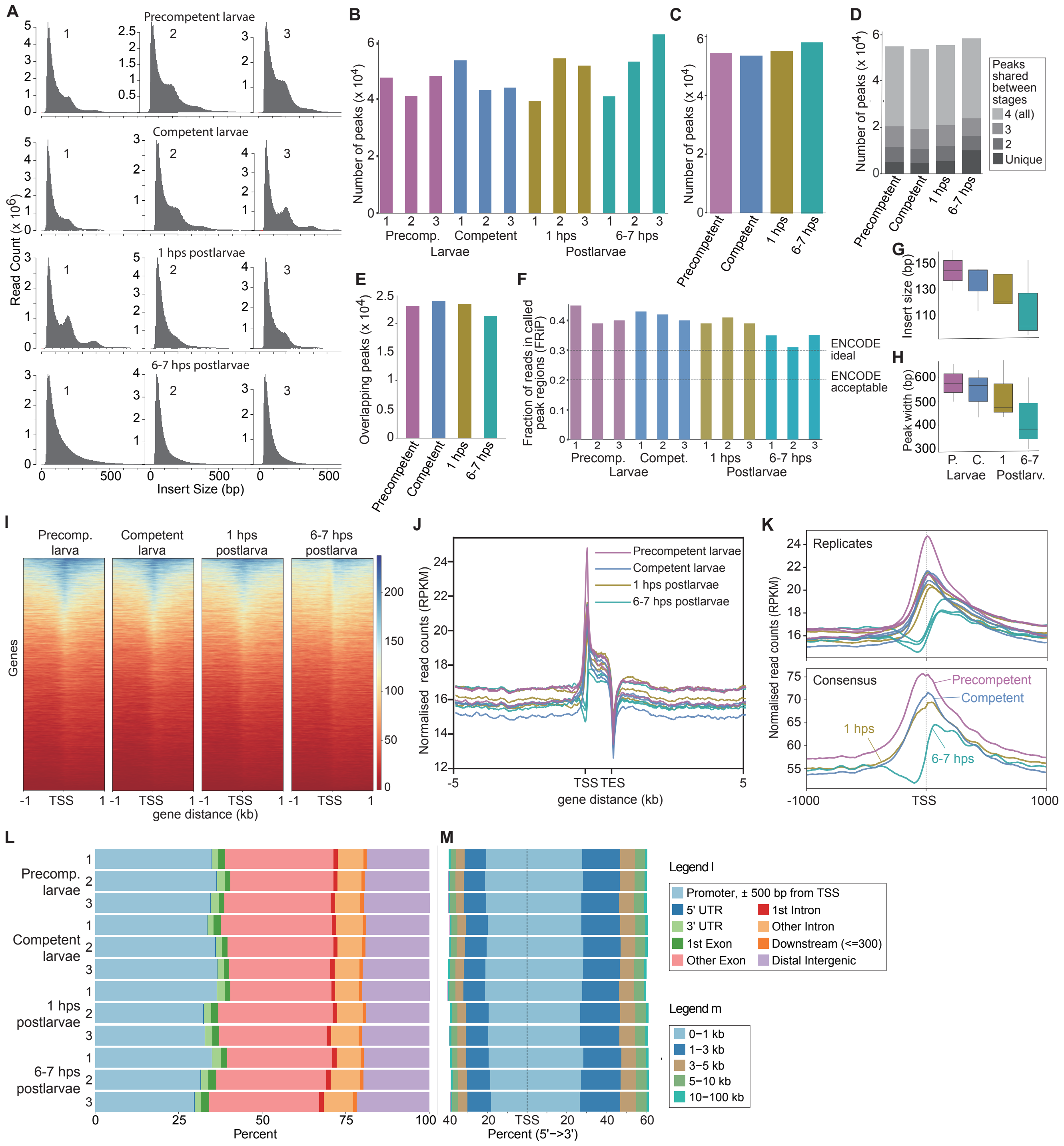

### FIg 3 supp 1

Supplementary Figure 3

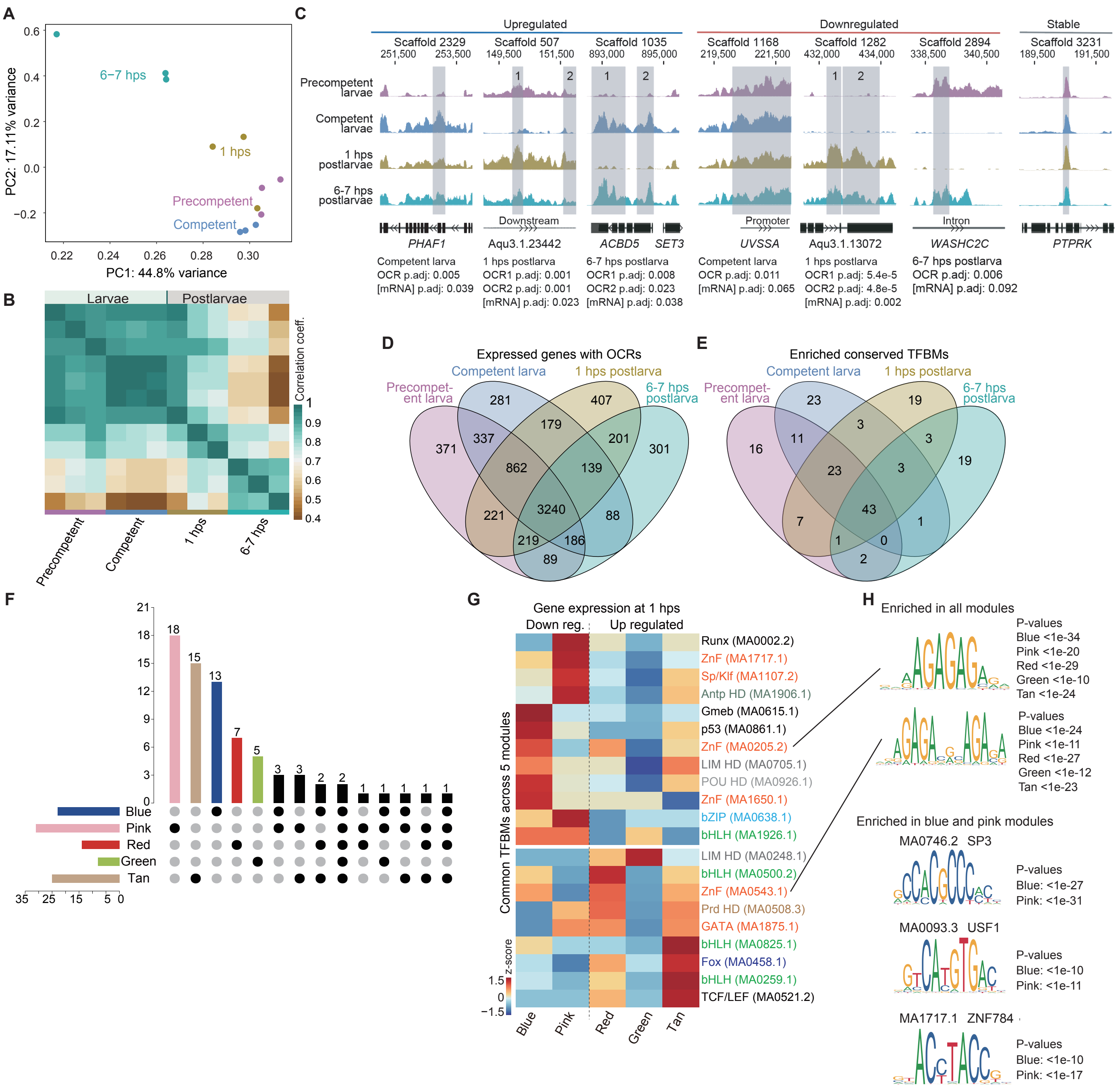
